# Systematic identification of pleiotropic genes from genetic interactions

**DOI:** 10.1101/112326

**Authors:** Elizabeth N. Koch, Michael Costanzo, Raamesh Deshpande, Brenda Andrews, Charles Boone, Chad L. Myers

## Abstract

Modular structures in biological networks are ubiquitous and well-described, yet this organization does not capture the complexity of genes individually influencing many modules. Pleiotropy, the phenomenon of a single genetic locus with multiple phenotypic effects, has previously been measured according to many definitions, which typically count phenotypes associated with genes. We take the perspective that, because genes work in complex and interconnected modules, pleiotropy can be treated as a network-derived characteristic. Here, we use the complete network of yeast genetic interactions (GI) to measure pleiotropy of nearly 2700 essential and nonessential genes. Our method uses frequent item set mining to discover GI modules, annotates them with high-level processes, and uses entropy to measure the functional spread of each gene’s set of containing modules. We classify genes whose modules indicate broad functional influence as having high pleiotropy, while genes with focused functional influence have low pleiotropy. These pleiotropy classes differed in a number of ways: high-pleiotropy genes have comparatively higher expression variance, higher protein abundance, more domains, and higher copy number, while low pleiotropy genes are more likely to be in protein complexes and have many curated phenotypes. We discuss the implications of these results regarding the nature and evolution of pleiotropy.

## Introduction

### Organization of functions in biological systems

Modularity of cellular functions has become a central tenet of systems biology, supported by evidence from diverse types of genomic data. Segal et al. (2003) designed a method that, from yeast gene expression data, inferred functionally coherent sets of genes that were regulated as a group according to different conditions. Gavin et al. (2006) described protein complexes in terms of core components and attached modules, using various data as evidence that grouped proteins act as single functional units. Costanzo et al. (2010) noted that the yeast genetic interaction (GI) network is well suited to define clustering of genes at various levels, from broad high-level biological processes down to specific pathways. With an eye to evolution, Ryan et al. (2013) found that most *S. cerevisiae* protein complexes are composed of genes that are either all essential or all nonessential. Further, this study noted that complexes conserved in *S. pombe* had the same property, but notably, proteins in some complexes switched essentiality as a group, indicating that this organization is favorable even in the context of evolutionary changes. Finally, Roguev et al. (2008) compared genetic interactions between the same yeast species and found evidence that while GIs are highly conserved within modules, a lower conservation of GI between modules allows “rewiring” to occur as the species diverge.

Despite the seemingly tidy nature of modules with these properties, considerable complexity characterizes modular organization due to substantial reuse and diverse effects of cellular components. Pleiotropy, when considered at the molecular level of genes, is the case in which perturbation of one gene influences multiple phenotypic traits (Paaby and Rockman, 2013; Stearns, 2010). For example, specific subcomplexes of nucleopores play important roles in gene silencing and DNA damage repair in addition to controlling nuclear import and export (Strambio-De-Castillia et al., 2010). As another example, multiple proteins responsible for mRNA decay in the cytoplasm, such as XRN1p, have a complementary chromatin-binding function that promotes genome-wide transcription initiation and elongation, mechanistically maintaining steady state mRNA levels (Haimovich et al., 2013). Famously, mammalian apoptosis pathways are triggered by components of the electron transport chain, such as cytochrome C (Ow et al., 2008). Other genes have a single molecular function but are fundamentally upstream of diverse cellular pathways, such as the HSP90 family of chaperones, which aid the folding of functionally diverse proteins (Taipale et al., 2010), and class V myosins, which use the actin cytoskeleton to localize mRNA and various organelles with help from cargo-specific receptor proteins (Hammer and Sellers, 2011). Because of the diverse physical interactors of the protein products, varied phenotypic effects appear when these genes are mutated.

In exploring the general notion of pleiotropy, researchers have used distinct definitions and datasets, showing that pleiotropy exists as many types of one-to-many genotype-to-phenotype relationships (Paaby and Rockman, 2013). All levels of biological organization have been considered: pleiotropy can connect DNA mutations or genes to phenotypic traits of molecular networks, cellular structures, organisms, populations, etc. Further, a phenotype may be described in the context of an environment, such as a genetic background, a population, a chemical, or nutrient availability. In humans, pleiotropy was recently explored by Pickrell et al. (2016), who used GWAS results to identify 341 loci in humans that are associated with multiple traits, including diseases. In yeast, phenotypic effects that stem from one gene have previously been measured by reverse genetics methods that screened the yeast deletion collection for phenotypes, such as measuring over 250 morphological phenotypes (Ohya et al., 2005) or measuring sensitivities to different stresses (Dudley et al., 2005; Ericson et al., 2006; Hillenmeyer et al., 2008, respectively assessing 21, 6, and 180 evironments). These studies variously estimate that between 5% and 30% of yeast genes could be considered pleiotropic according to counted numbers of traits or environmental sensitivities. Although different environmental challenges can require different functional roles, these studies do not link conditions to specific functions, leaving the possibility that genes sensitive to many environments may belong in a generalized stress response category.

Given the extensive sets of gene annotations assembled by The Gene Ontology (Ashburner et al., 2000) for human and model organisms, counting annotations is a natural way to identify pleiotropic genes and has been employed in a number of studies. Khan et al. (2014) used semantic similarity of GO terms that clustered into non-overlapping functions to identify moonlighting proteins, a strict but particularly interesting type of pleiotropy in which functions are physically separable but not as a result of physical partitioning in the protein. The authors found that moonlighting proteins often (48% of the time) contain disordered regions. Pritykin et al. (2015) carefully considered the structure of the GO tree in order to identify genes with distinct functions. The multifunctional genes were tested for associations with a number of gene properties, revealing that multifunctional genes are more likely to be large and multi-domain, essential, broadly expressed, central in PPI networks, have many regulators, and contain disordered regions.

### A genome-wide and modular basis for pleiotropy

Biological networks, as structures, can naturally represent modularity, redundancy, and pleiotropy, providing detailed context that allows more comprehensive understanding of cell function (Kim and Przytycka, 2012; Vidal et al., 2011; Yu et al., 2016). In fact, gene functions are not limited to associations with visible phenotypes: genes can affect network properties, such as support robustness and flexibility (Burga et al., 2011; Levy and Siegal, 2008; Park and Lehner, 2013; Rutherford and Lindquist, 1998). Therefore, estimating pleiotropy at a molecular level as effects measured within a network is of particular importance.

In protein interaction networks, a popular network-based characterization of hub proteins is classification as an intra-modular (“party”) node, which mainly functions as part of a module and has correlated expression with its neighbors, or an inter-modular (“date”) node, which coordinates between modules or has multiple functions (Agarwal et al., 2010; Han et al., 2004; Pritykin and Singh, 2013). A strength of physical protein interactions is that they are mechanistically interpretable; however this type of relationship is limited by physical locality. In contrast, genetic interactions identify a variety of functional relationships, including partial redundancy within the same module, pathway buffering, and dependency/similarity within a spatially or temporally directed pathway. We believe that genetic interactions provide a novel and valuable view of pleiotropy because they (i) are known to appear both within and between pathways, (ii) capture functional relationships between genes that operate in different high-level processes, (iii) can recover functions that are buffered by other genes, and (iv) can reflect any biological process, not just those in a restricted set of measured phenotypes. This last point is a solution to the problem of trait selection that many estimates of pleiotropy are bound by. The GI network is therefore an informative place to assess the molecular pleiotropy of genes.

In this work, using genetic interactions, we consider pleiotropy to be one gene affecting multiple sectors of cellular function such that there is a phenotypic consequence of fitness defect. With this definition, we are able to characterize the properties and behavior of genes that impinge on diverse functional modules and affect multiple traits at the molecular level.

For measuring pleiotropy in this study, we extract modules from the GI network using a data mining method originally described by Bellay et al. (2011), who used frequent item set mining to exhaustively discover modules of genetic interactions, termed biclusters, that covered the majority of the yeast genetic interaction network (Costanzo et al., 2010). A bicluster is composed of two sets of genes and each gene in one set interacts with every gene in the other set—put another way, it is a complete bipartite subgraph of the GI network. Biclusters represent genes with similar behavior, because all genes on one side of a bicluster share a set of interaction partners; in this way, they are similar to clustered gene profiles, a popular framework for identifying functionally related genes. However, biclusters are built from subsets of a gene’s interaction partners, meaning they can identify multiple functions per gene and thus represent reuse in addition to modularity. Bellay et al. (2011) found that the bicluster coverage of interactions in a hub gene’s profile may relate to pleiotropy, since this correlated with the number of GO terms annotated to the gene as well at the number of drug sensitivities (Hillenmeyer et al., 2008). In the following analysis, we describe a novel definition of pleiotropy derived from GI biclusters in a new, nearly complete yeast GI network (Costanzo et al., 2016). We measure pleiotropy using an entropy measure computed on the set of gene’s genetic interaction biclusters to describe the functional spread of a gene’s effects on phenotype. We evaluate characteristics of the high- and low-pleiotropy genes identified by our approach and report several physical, functional, and evolutionary properties that differ between the two pleiotropy classes.

## Results

### Measuring pleiotropy from participation in GI modules

Genetic interactions capture relationships between genes involved in different processes, and biclusters, which are groups of genes densely connected by genetic interactions, are able to characterize the functional context of these relationships. We define a measure of pleiotropy that expresses a gene’s functional distribution within these bicluster modules (Figure 1). Our first step was to apply XMOD to an input GI network to obtain its set of biclusters. For each gene, we collected all biclusters that contain it and removed clusters that were redundant (see Methods). A bicluster consists of two sets of genes, densely connected by a set of genetic interactions bridging them. In the context of calculating pleiotropy for a specific gene *G*, we refer to the set containing *G* as the “associate side” and the set of genes interacting with *G* as the “adjacent side.” We use simple criteria to annotate biclusters with high level biological processes: if the associate-side genes are statistically enriched for or are at least 40% composed of genes annotated by a term, then the bicluster is labeled with that term. Having identified and annotated a gene’s biclusters, we then count the process annotations, resulting in a functional profile of the gene’s modules (Figure 1, 2A). The final pleiotropy score is the entropy of the process annotations counted in the profile (Figure 1, 2A). Entropy is a non-negative value that is 0 in the case that all annotations are the same and reaches a maximum when all possible annotations occur an equal number of times. The number of terms used for annotations, not the number of annotations a gene’s biclusters receive, determines the maximum value entropy can reach. We used a set of 20 manually annotated (MA) biological processes and the entropy scores range between 1 and 4 (Figure 2B). See Methods for more details on our measurement of pleiotropy.

**Figure 1.**
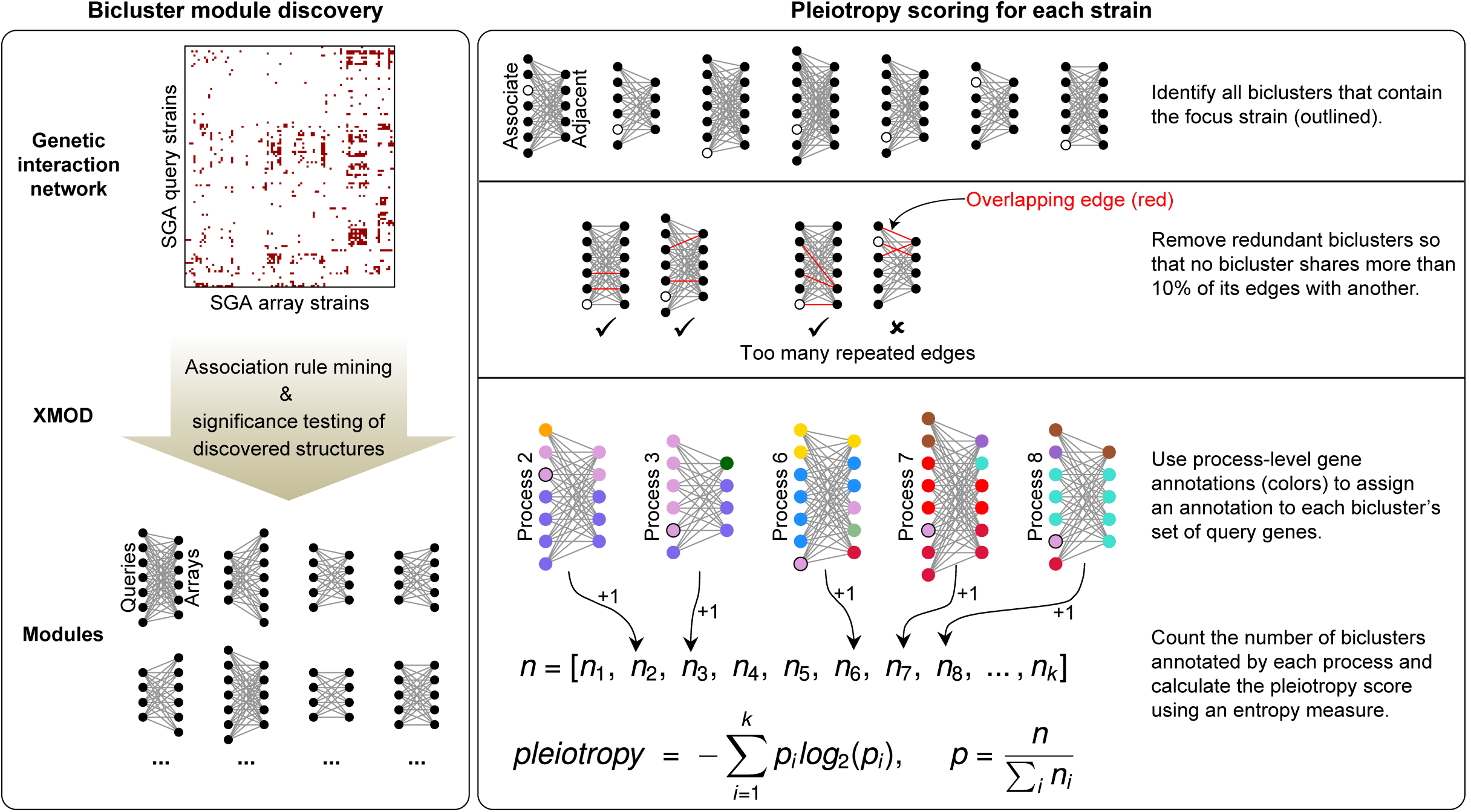
Measuring pleiotropy from GI modules. Bicluster modules are obtained by applying XMOD to the genetic interaction network (Left box). The input SGA-derived network is binarized by thresholding negative genetic interactions between query and array single mutant strains, reflecting the SGA experimental setup. Interactions are added between query and array strains representing the same gene to allow modules containing these. Discovered complete bipartite modules have one set of query strains and one set of array strains. The pleiotropy of a focal strain, depicted as an outlined circle, is calculated from the functional distribution of its containing bicluster modules (Right box). Bicluster annotations are determined by the associate side of the module, the set of genes that contains the focal gene and is drawn as the left side of each bicluster (the right strain set is the adjacent side). Colors represent gene annotations. The vector **n** contains counts of annotation occurrences.

**Figure 2.**
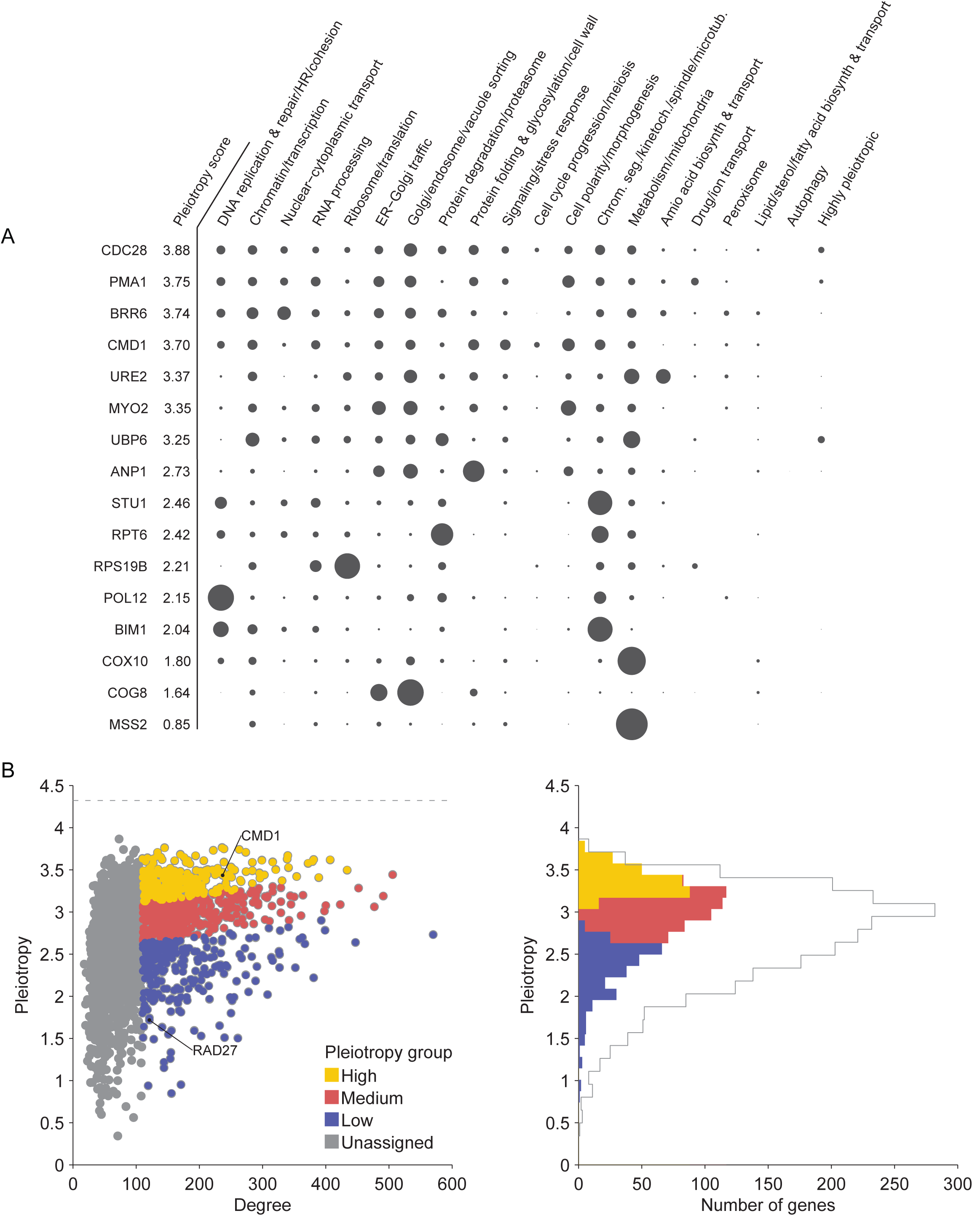
Pleiotropy scores. (A) Functional profiles of example genes with a range of pleiotropy scores are sorted with the most pleiotropic at the top. The distribution of annotation occurrences from each gene’s containing modules was normalized (equal to vector **p** in Figure 1) and displayed such that circle area represents the fraction of module annotations in each bioprocess. Data for each are from a single query strain. (B) Gene pleiotropy scores are correlated with genetic interaction degree, but still show substantial variation not explained by degree. High, low, and medium pleiotropy groups are only assigned to genes with degree of at least the 60^th^ percentile (vertical boundary between gray and colored markers, left plot) and are determined from residuals of the regression of pleiotropy scores against degree (cause for sloped divisions between the high-medium and medium-low boundaries, left plot). Histogram bars in the right-hand plot are stacked and count the genes assigned to pleiotropy groups. Both plots show DMA-derived pleiotropy scores.

In implementing this pleiotropy measure, we used negative genetic interactions of the latest, near-complete yeast GI network (Costanzo et al., 2016). This network comprises two distinct datasets reflecting the experimental organization used in its construction. The separation of the two GI networks persists throughout our work here: we derived pleiotropy scores from each. The first GI network, called the TSA (temperature sensitive array) network, contains 2112 query genes screened for interactions with 560 essential array genes and 173 nonessential array genes. The second, called the DMA (deletion mutant array) network, contains 4004 query genes screened for interactions with 3827 nonessential array genes. The query genes of both networks include nonessential genes, experimentally represented by gene deletions, and essential genes, which were represented by temperature sensitive and DAmP alleles. Accordingly, the biclusters from both networks can have a mixture of essential and nonessential genes on one side, the query side. We focused primarily on measuring pleiotropy for query genes (more precisely strains, see Methods), so in this case the associate-side module enrichment step of our pleiotropy method analyzed mixed-essentiality groups of genes. We also implemented our pleiotropy measure with a different data orientation, computing pleiotropy scores for array instead of query genes, and with a second annotation scheme, the experimentally derived SAFE annotations instead of the manual set. We use the term “scoring configuration” to refer to a data setup used in generating pleiotropy scores, which specifies the GI network, annotations, and type of strains analyzed; in total, there are six configurations, which are described in the Methods.

When using the genetic interaction network, a straightforward pleiotropy metric could be the number of interactions observed for a given gene. A gene’s genetic interaction degree is very informative about the magnitude of a mutation’s effect. For example, degree is strongly correlated with fitness defect (Pearson’s r = 0.78, p < 10e-300; (Costanzo et al., 2016, nonessential strains)), and is also correlated with the number of GO terms (r = 0.23, p < 10e-42) and the number of curated phenotypes (r = 0.65, p < 10e-300), two gene features that can indicate multiple functions. The pleiotropy score we developed, however, is more specific than GI degree—it is designed to distinguish different functions of a gene, first by organizing GIs into modules, and second, by assessing annotation profiles with fractions instead of counts in the entropy calculation. This decoupling of degree and pleiotropy is evident by the variation depicted in Figure 2B, which illustrates how a high degree alone is not sufficient for a gene to have high pleiotropy. Nevertheless, the Spearman correlation of 0.45 (p < 10e-53) between entropy and degree suggests that attempts to characterize pleiotropic genes may simply recover trends already associated with degree. To focus specifically on the functional breadth of genes independent of their interaction degree, we controlled for GI degree when defining pleiotropy classes. Specifically, we first take pleiotropy as the residual of the regression of entropy against degree. We then limit genes to those that have a negative GI degree at or above the 60^th^ percentile. Finally, we classify these high-degree genes as high, medium, or low pleiotropy by binning scores into the highest 30%, middle 40%, and lowest 30%. These three classes are used for all statistical analyses discussed in the following sections. As previously mentioned, we used the TS-derived and DMA-derived GI networks separately in measuring pleiotropy, therefore we have a set of three pleiotropy classes for each network. We specify source GI data in the text when discussing specific results.

### Many primary functions are represented in high-pleiotropy genes

Many of the genes that displayed high pleiotropy have known associations with particular functional pathways. The chaperone HSP90, whose pleiotropy score is in the highest 30%, is a classic example of how participation in a central maintenance pathway allows the gene to suppress phenotypic variation in many aspects of cellular biology. We found that this is not unique; genes involved in many other cellular functions also exhibited high pleiotropy. The following are brief examples of some of the many functional annotations already associated with genes in our high pleiotropy class: cell cycle regulation (CDC28, CKS1, cyclin CLN3, whole genome duplicates SWI5 and ACE2, RAM pathway component TAO3); the ubiquitin system (UBI4, UBP1, DOA1, CDC53, RAD6, RSP5, TOM1, UBP6, UBR2, HRT1, UFD1, UBP14); stress response and protein folding (chaperones HSP82, CDC37, and CNS1, HSP82 regulator HSP1); membership in the CCR4-NOT complex, a global transcription regulator (CDC36, CDC39, NOT3, CAF120); ribosome biogenesis (MAK11, NEW1, DBP7); nuclear-envelope membrane functions (BRL1, BRR6, and APQ12); and vacuole functions (VPS62, VAC7, VAC14, VPS66, ZRT3, IML1).

### Calmodulin’s biclusters illustrate broad functional influence

As another example, we highlight the high-scoring pleiotropic gene CMD1 (Figure 3A, Figure 2A), which encodes the binding protein calmodulin and is conserved in all eukaryotes. It is well known to regulate many processes, a functional ubiquity that likely is enabled mechanistically by the capacity to bind calcium ions in four different sites in most species as well as target proteins, many of which trigger function-specific conformations of calmodulin. Evidence of binding site functional specificity comes from Ohya and Botstein (1994), who found four groups of mutations that resulted in distinct phenotypes. Using its namesake ability to detect Ca+ ions, CMD1 activates calcineurin and two protein kinases when Ca2+ concentration is high, which control a number of downstream processes (Cyert, 2001). Within the GI network, CMD1 appears in dozens of biclusters. Nine of them are shown in Figure 3A to illustrate both how GI-derived pleiotropy is apparent from structured modules and the functional coherency that characterizes these modules. One of Calmodulin’s known localizations is the bud tip and neck due to its physical interaction with MYO2p, a myosin protein that is required for polarized growth (Stevens and Davis, 1998). This relationship is reflected in the bicluster labeled “Cell polarity/morphogenesis”, which contains cell wall integrity genes SLT2 and BCK1, bud neck and wall localized proteins SKT5, CHS3 (recruited by SKT5 and MYO2), and ROM2, ARP2/3 activator PAN1, and polarity-establishing BEM1 (Duncan et al., 2001; Levin, 2005; Madden and Snyder, 1998). Another established localization behavior of calmodulin is association with the spindle pole body throughout the cell cycle. During mitosis, it is involved in attachment of microtubules to the SPB and is required for correct spindle function (Sundberg et al., 1996). This explains its membership in the bicluster labeled “Chrom. seg/kinetoch./etc” along with spindle organizers CIK1 and STU2 (a SPB-interactor), as well as a number of kinetochore genes, AME1, OKP1, and NSL1, and kinetochore recruitment gene CTF13 (De Wulf et al., 2003; Kosco et al., 2001; Page et al., 1994). The shared negative interactors of these genes, adjacent in the bicluster, are genes from the spindle assembly checkpoint (SAC), which can buffer a dysfunctional spindle by prolonging prometaphase. Lastly, Cmd1p is thought to regulate the final stages of vacuolar fusion (Peters and Mayer, 1998). The bicluster labeled “Golgi/endosome/vacuole” reflects this role, containing two components of the cytoplasm-to-vacuole targeting pathway complex TRAPPIII and GYP1, which respectively activate and deactivate the vesicle docking regulator YPT1, as well as SEC17, which is required before vacuole membrane fusion events, and three members of the COG complex (Du and Novick, 2001; Lynch-Day et al., 2010; Ungermann et al., 1998). A short discussion of some of the remaining modules in Figure 3A is in the supplement. Though GI modules do not explain specific functions of a gene, the example of CMD1 shows how genetic interactions can recover evidence for functions established in the literature.

**Figure 3.**
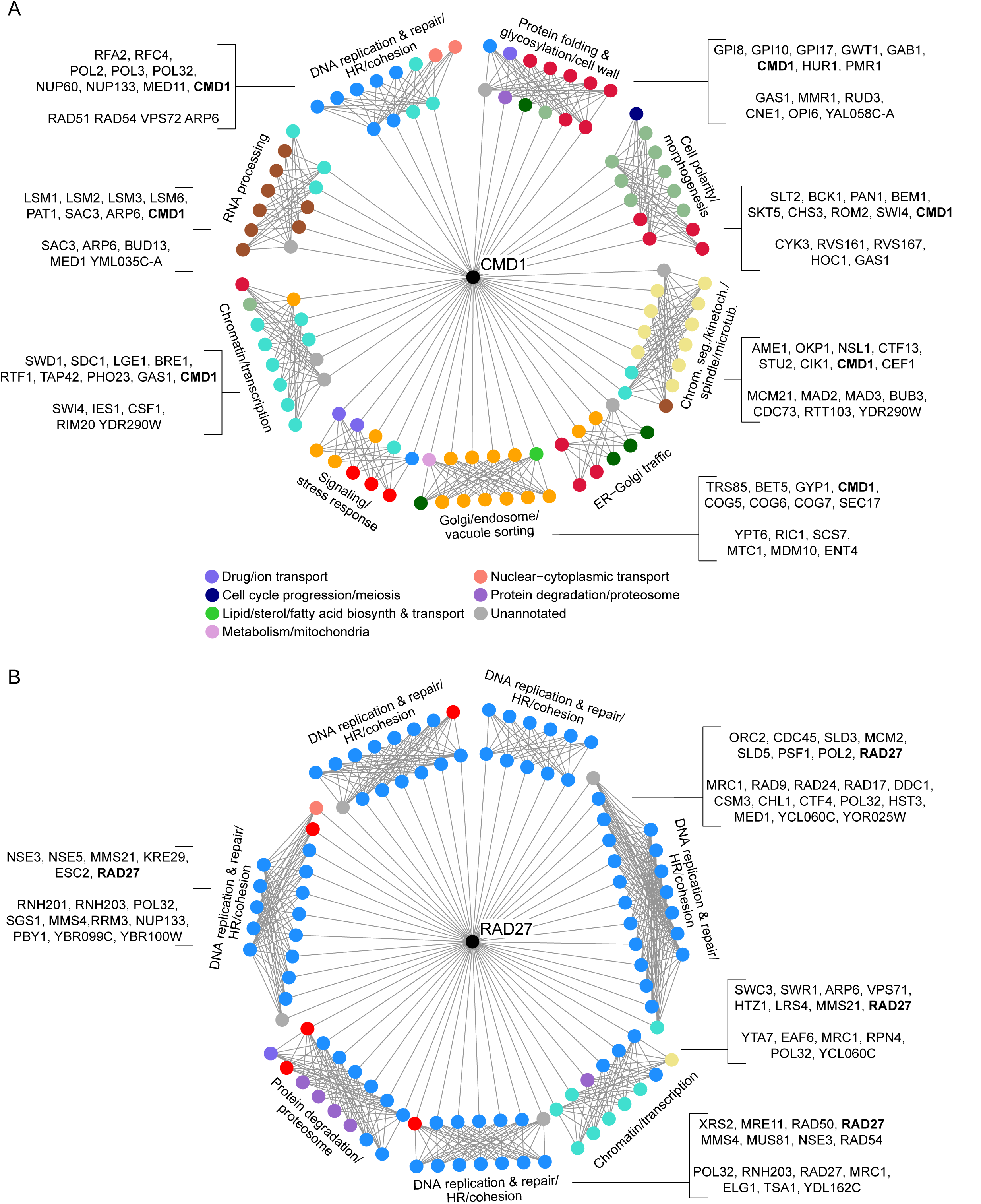
Selected biclusters of the pleiotropic gene CMD1 (A) and the non-pleiotropic gene RAD27 (B). Nodes represent genes and edges represent negative genetic interactions extracted from the DMA-derived GI network. Only genetic interactions that define each bicluster are displayed, although there are often interactions between genes on the same side of a bicluster. The biclusters’ adjacent-side sets of genes are connected to the focal genes CMD1 or RAD27. The genes arranged on the outside of the each diagram, with the addition of the focal gene, are the associate sides. Gene names list the members of some biclusters; the first group of names in each bracketed pair lists the associate-side genes and the second group lists the adjacent-side genes. Text labels are bicluster annotations determined from the associate-side genes. Colors of nodes indicate the functional annotations of genes, which can be inferred from the bicluster annotations (*e.g.* sea green represents “Cell polarity/morphogenesis”). Any colors that cannot be interpreted with bicluster labels are listed in the legend. Any genes that have multiple process annotations are colored preferentially to match the annotation given to the bicluster. Both panels use the same color scheme.

In contrast to the highly varied functions of CMD1, we also make an example of RAD27 (Figure 3B). This gene has a focused functional influence on cellular processes, and therefore low pleiotropy, with nearly all of its containing biclusters representing DNA replication and repair functions. Despite the clear theme of RAD27’s modules, we do see individual pathways clustering together. For example, the associate side of one bicluster contains genes in complexes that initiate and drive the replication fork during DNA replication (Medagli et al., 2016). The genes PSF1 and SLD5, as half of the GINs complex, and SLD3 help to assemble the pre-inititation complex, which includes MCM2, ORC2, and CDC42, at replication origin sites. Many of these genes go on to form the CMG complex, the helicase that unwinds duplex DNA and progresses in the core of the replication fork. This set of genes negatively interacts with genes involved mitotic checkpoints for DNA damage and DNA replication, MRC1, RAD9, RAD24, DDC1, RAD17, and CSM3, which appear in the bicluster’s adjacent side. Another of RAD27’s biclusters contains histone-related genes in both sides (e.g. SWC3, SWR1, ARP6, VPS71, HTZ1, YTA7, and EAF6), and others contain a number of genes related to RAD27’s known functions, Okazaki fragment processing and double-strand break repair (e.g. POL31, RNH203, RNH201, XRS2, MRE11, and RAD50).

### GO term enrichment within pleiotropy classes

In order to discover any particular cellular processes or components that are significantly biased in their composition of pleiotropic genes, we performed hypergeometric tests for enrichment of GO terms in our high and low pleiotropy classes. We found that high pleiotropy genes were not enriched for any GO terms. Although this result is somewhat surprising, it is consistent with the observation that pleiotropic genes work in diverse primary functions. Low-pleiotropy gene classes from both GI networks were enriched for a number of terms. Low-pleiotropy genes derived from the DMA network were enriched for golgi vesicle-mediated transport, as well as more general transport and localization terms, and mitochondrial respiration. For example, 43 of all 55 background genes annotated by the GO component “mitochondrial inner membrane” have low pleiotropy (enrichment, p < 10e-11). The low-pleiotropy class derived from the TSA network was enriched for vesicle transport also, and DNA replication and proteolysis terms. For example, 15 of the 16 genes in the cytosolic proteasome complex have low pleiotropy (enrichment, p < 10e-4).

### Characterizing high- and low-pleiotropy genes

Next, we searched for evolutionary and functional characteristics of high- and low-pleiotropy genes by testing for associations with 37 gene and protein properties. A number of gene properties differed significantly between the two pleiotropy classes in Wilcoxon ranksum tests (Figure 4, Table 1). High-pleiotropy genes were positively associated with expression variation, high gene copy-number-based features, high protein abundance, and many domains, while the low-pleiotropy genes tended to participate in protein complexes and, surprisingly, had more curated phenotypes. Specific statistics presented in the text below are based on pleiotropy scores of query strains, our default analysis set, derived from both the TS and DMA GI networks. Our reporting of results is conservative: we tested 22 variations of our method and report results that are robust across many (Methods and Table 1). For example, SGA interrogates essential genes with mutant strains that are temperature-sensitive point mutations or DAmP (low expression) alleles. Because it is easy to imagine a point mutation that affects only the subset of a gene’s functions that is dependent on a single part of the protein, one variation of our ranksum tests excludes TS strains, leaving just DAmP alleles to represent the behavior of essential genes. An extended description of all testing variations performed and the set of results for each scoring configuration are presented in the supplement.

**Figure 4.**
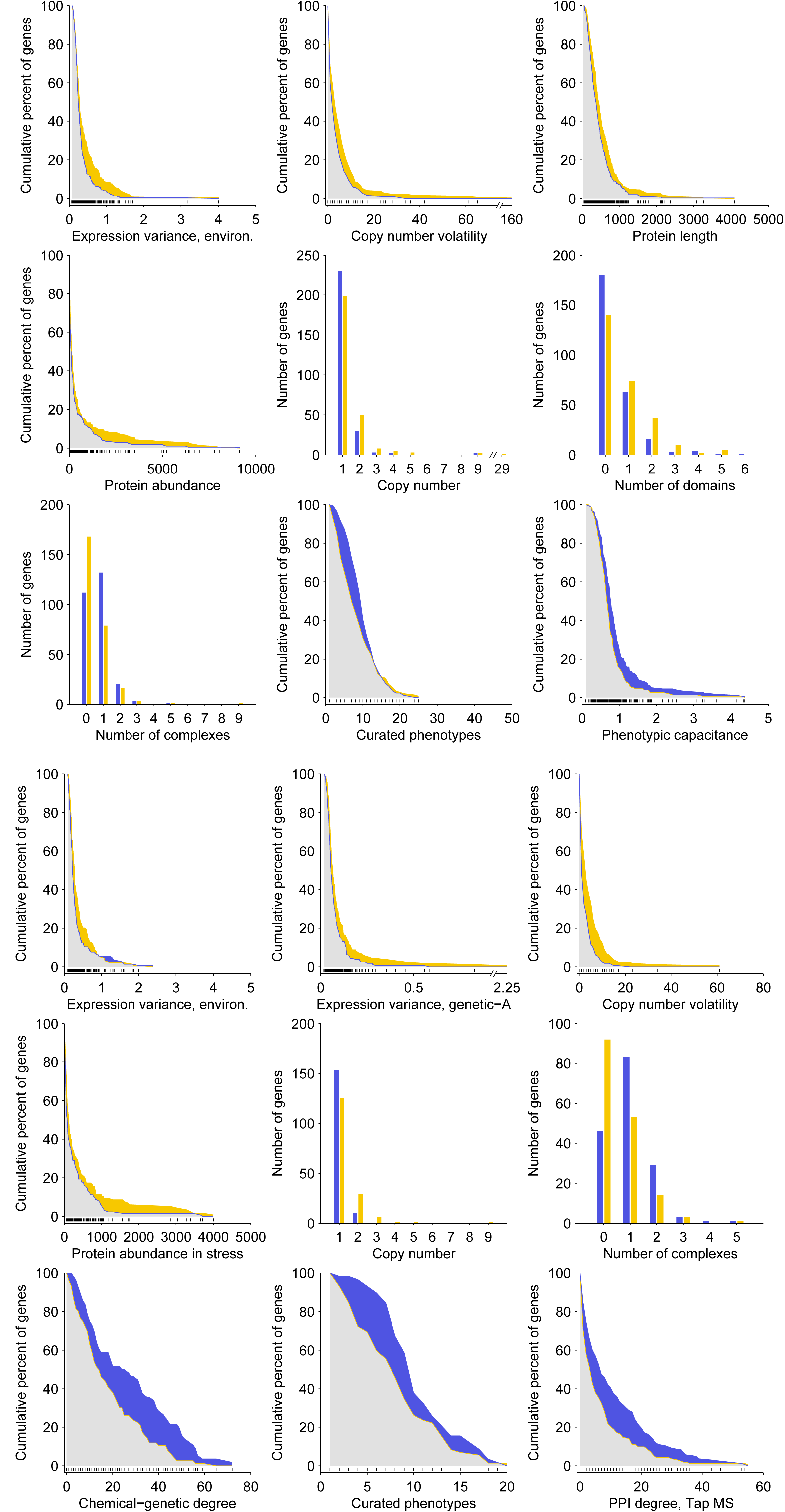
Gene properties significantly associated with high (yellow) or low (blue) pleiotropy derived from the DMA (A) and TSA (B) GI networks. Cumulative plots are displayed for properties that take on many values. For a pleiotropy group and property, the plotted line shows the percent of the genes that have a property value greater than or equal to any point on the x-axis. Percentage calculations only take into account genes that have measured values for the property (all properties shown had sufficient data coverage, see Methods). The area between the blue and yellow lines is filled with color indicating which pleiotropy group has a higher percent of genes with high property values. Black hash marks plotted above the x-axis mark all values found in the genes’ property values. Bar plots are displayed for properties that take on few values. There is a total of 268 genes in both the high and low DMA-derived groups (A) and a total of 163 genes in both TSA-derived groups (B). P-values from ranksum tests for each property from left to right are (A, top row) 7e-3, 8.4e-4, 7.4e-4; (A, second row) 3.4e-2, 4.3e-4, 1.1e-4; (A, third row) 1.1e-5, 2.7e-4, 1.6e-3 (B, top row) 5.4e-3, 6.4e-3, 2.4e-6; (B, second row) 3.4e-2; 8.7e-6, 7.5e-7; (B, third row) 5.8e-3, 2.1e-3, 5.9e-5.

**Table 1.** Summary of gene characteristics associated with high- and low-pleiotropy genes. Tests were performed for pleiotropy scores derived from different pleiotropy scoring configurations (columns). Values shown are the number of ranksum tests that yielded a significant p-value, out of a total of 22 variations performed for each query-strain scoring configuration and 12 variations performed for each array-strain scoring configuration (see Methods). Blank cells indicate zero tests with significant results. Values in parentheses indicate significant results that contradict the result column by associating the gene property with the opposite pleiotropy class. Asterisks indicate features that were associated strongly enough with both pleiotropy classes that the property is listed in two rows. The significance of p-values from ranksum tests was determined using the FDR-control procedure described in Benjamini et al. (2006), counting tests for 37 gene properties as a family.

| Gene/protein property | Pleiotropy class with positive association | DMA |  |  |  |  |  | TSA |  |  |  |  |  |
| --- | --- | --- | --- | --- | --- | --- | --- | --- | --- | --- | --- | --- | --- |
|  |  | Query |  |  | Array |  |  | Query |  |  | Array |  |  |
|  |  | Associate, MA | Associate, SAFE | Adjacent, MA | Associate, MA | Associate, SAFE | Adjacent, MA | Associate, MA | Associate, SAFE | Adjacent, MA | Associate, MA | Associate, SAFE | Adjacent, MA |
| Expression variance, environ. | High | 9 | 16 | 10 | 3 | 5 | 6 | 17 | 22 | 22 | 2 | 3 | 2 |
| Protein abundance in stress | High | 3 | 16 | 21 | 10 | 12 | 1 | 9 | 14 | 22 |  |  | 3 |
| Protein abundance | High | 5 | 16 | 21 | 12 | 12 | 1 | 1 |  | 19 |  |  | 2 |
| Expression variance, genetic-A | High |  | 15 |  |  | 1 | 4 | 17 | 22 | 18 | 6 |  | 1 |
| CAI | High | 1 |  | 20 |  | 12 |  | 1 | 15 | 9 |  |  | 2 |
| Copy number | High | 22 | 18 |  | 4 | 8 |  | 22 | 22 | 22 |  |  |  |
| Copy number volatility | High | 22 | 18 |  | 12 | 8 |  | 22 | 22 | 15 |  |  |  |
| WGD duplicate | High | 21 | 18 |  | 3 | 8 |  | 21 | 22 | 16 |  |  |  |
| Expression variance, genetic-B | High |  |  | (L 15) | 5 | 1 | 4 | 1 | 18 |  | 2 |  |  |
| Single mutant fitness defect* | High |  |  | 11 |  | 5 | 6 |  |  |  | 2 | 12 | 8 |
| Transcription level | High |  |  | 18 | 5 | 5 |  |  | 1 | 19 |  |  | 3 |
| Expression level | High | 1 |  | 18 | 5 |  |  |  |  | 19 |  |  | 3 |
| Number of domains | High | 21 | 22 | 15 |  |  |  |  | 2 |  |  | 5 |  |
| Number of unique domains | High | 21 | 22 | 15 |  |  |  |  | 2 |  |  | 5 |  |
| Originated in Saccharomyces | High |  |  |  |  |  |  | 22 | 16 | 9 |  |  |  |
| Protein length | High | 22 | 19 | 5 |  |  |  |  |  |  |  |  |  |
| Coexpression degree | High |  |  | 10 |  |  |  |  |  | 10 |  |  |  |
| Essential* | High | 7 |  | 20 |  |  |  |  |  |  |  |  |  |
| Complex member | Low | 22 | 21 | 1 | 12 | 8 |  | 22 | 22 | 21 | 3 |  |  |
| Number of complexes | Low | 22 | 18 | 1 | 12 | 8 |  | 22 | 22 | 20 | 6 |  |  |
| Curated phenotypes | Low | 22 | 22 | (H 10) | 3 | 5 | (H 6) | 19 | 21 | 11 |  |  |  |
| Phenotypic capacitance | Low | 21 | 21 |  |  | 2 |  | 7 | 19 | 10 |  |  |  |
| Chemical-genetic degree | Low |  |  |  |  | 12 | 2 | 20 | 22 | 1 |  |  |  |
| Effective number of codons | Low |  |  | 5 | 9 | 10 |  |  |  | 21 |  |  | 1 |
| Multifunctionality | Low |  |  | (H 2) | 7 | 4 |  | 18 | 21 | 3 |  |  |  |
| PPI degree, Tap MS | Low |  |  | (H 4) | 3 |  |  | 21 | 4 | 5 | 7 |  |  |
| Single mutant fitness defect* | Low | 22 | 22 |  | 3 |  |  | 8 | 1 |  |  |  |  |
| Yeast conservation | Low |  | 3 | (H 2) | 1 |  |  | 11 | 20 |  |  |  |  |
| dN/dS | Low | 1 |  | 13 |  | 10 | 6 |  |  |  |  |  |  |
| Age 1-20 | Low |  |  |  |  |  |  | 19 | 6 | 5 |  |  |  |
| Broad conservation | Low |  |  | (H 2) |  |  |  | 17 | 1 |  | 2 |  |  |
| Essential* | Low |  |  |  |  |  |  | 19 | 10 | 15 |  |  |  |

### Expression variance and protein abundance are higher among genes with broad influence

Two different measurements of gene expression level variance were robustly associated with high-pleiotropy genes. Environmental expression variance is determined by subjecting yeast cells to many environments and measuring gene expression levels, then calculating variance for each gene (Gasch et al., 2000). A Wilcoxon ranksum test showed that genes in the high pleiotropy class had higher environmental expression variance than genes in the low-pleiotropy class, using both the DMA (p < 7e-3) and TSA (p < 6e-3) pleiotropy scores (Expression variance, environ., Figure 4A,B). Among the 50 genes with the highest environmental expression variance, 22 have high DMA-derived pleiotropy compared to only 5 with low pleiotropy (Figure 5A; 23 have a medium pleiotropy). We found that, regardless of pleiotropy level, most genes with high variance reached their extreme expression levels during heat shock and cold shock conditions and during stationary phase. Regulatory response to environmental stresses consists of induced expression of some genes and suppression of others, a program that is similar in all stress environments, not condition-specific (Gasch et al., 2000). We find that there is no bias in the high pleiotropy genes towards having increased or decreased expression during stress conditions (Figure 5B).

**Figure 5.**
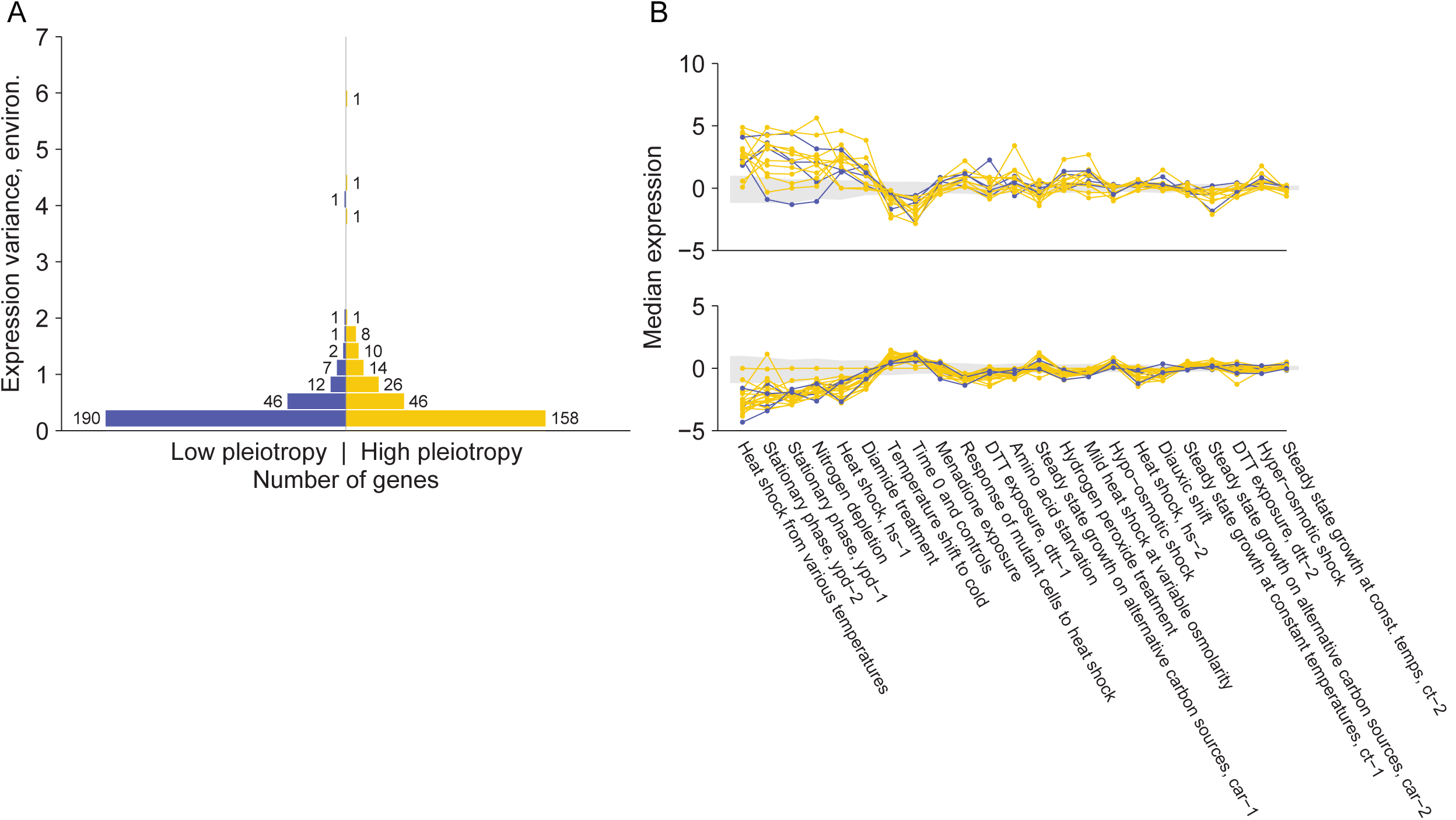
High-pleiotropy genes have higher environmental expression variance than low-pleiotropy genes. (A) Histograms show the distribution of expression variance, shown on the y-axis, for all high- and low-entropy genes. The number of genes in each pleiotropy class that fall into each bin is given next to each bar. (B) Expression of all high- and low-pleiotropy genes that are among the 60 genes (regardless of pleiotropy) with the highest expression variance was plotted for each environment. Values plotted are medians of multiple time points measured for individual environments. Separate axes are for visual clarity only: genes that tend to increase in expression during stress are plotted on the top, while genes with the opposite trend are on the bottom.

Genetic expression variance, calculated from the genome-wide expression profiles of many crosses between the diverged *S. cerevisiae* strains BY and RM (Expression variance, genetic-A, Figure 4B), was also associated with high pleiotropy genes identified from both GI networks (DMA-derived pleiotropy, SAFE annotations: p < 7e-3; TSA-derived pleiotropy: p < 7e-3), although this result depended on the configuration used for the DMA-derived pleiotropy classes (Table 1, Supplement). Among the 50 genes with the highest variance, 22 had high pleiotropy and 7 had low pleiotropy (21 have medium pleiotropy). Similarly, a second measure of genetic expression variance (Expression variance, genetic-B, Table 1), which is measured from the gene expression of many diverged and geographically varied *S. cerevisiae* strains, was associated with high pleiotropy genes for a number of scoring configurations. For one scoring configuration, in which pleiotropy of DMA queries was measured using adjacent-side bicluster enrichments, this expression variance feature was associated with low-pleiotropy genes.

The two expression variance measures strongly associated with high-pleiotropy genes, environmental and genetic-A, had a Pearson’s correlation of 0.21 (p < 2.6e-13), suggesting that highly variable genes defined by the two measures overlap. However, environmental variance remained robustly associated with pleiotropy after controlling for the genetic-based feature (TSA-derived pleiotropy, ranksum p < 0.017). Genetic expression variance was not correlated after controlling for the environmental feature, suggesting that environment-induced expression variation is more strongly linked with high pleiotropy.

Protein abundance levels offer further observation of cellular usage of a gene, since translation and protein degradation are regulated. Protein abundance, including protein abundance under stress conditions, tended to be higher in high-pleiotropy genes than in low-pleiotropy genes (nonstress, SAFE p < 7e-3; stress, SAFE p < 5e-3; Figure 4).

### Copy number is higher in high pleiotropy genes

High-pleiotropy genes tended to have higher copy number, which is the number of genes that evolved from a single ancestral gene (paralogs) of the focal gene, compared to low-pleiotropy genes (DMA, SAFE p < 2e-3; Figure 4A, B). High pleiotropy genes with a copy number greater than two are in protein families that function in environmental responses as transmembrane proteins or components of signaling pathways, consistent with previous characterization of genes that have frequently duplicated (Wapinski et al., 2007). The most extreme copy number is that of high-pleiotropy gene RGT2, which, with its paralogs, is in a family of transmembrane sugar-transport channels, including some that trigger response to intracellular sugar concentrations. A more well-known example is the hub IRA2, which is a negative regulator of RAS2 and has two paralogs.

Duplicate gene pairs that arose from an ancient *S. cerevisiae* whole-genome duplication (WGD) event are distinguished from all other duplicate pairs, which resulted from small-scale duplication (SSD) events (Guan et al., 2007; Hakes et al., 2007; Kellis et al., 2004; Wapinski et al., 2007; Wong et al., 2002). We found that this difference is important with respect to pleiotropy. WGD genes were strongly associated with high pleiotropy genes, while SSD genes had only slight evidence of an association (Table 1, full table in supplement). This difference is difficult to explain, since there is no consensus on how evolutionary models apply differently to these scenarios. However, it is possible the genome state after a whole-genome duplication helps genes diversify by providing broader redundancy, like entirely duplicated complexes and pathways.

The DMA-derived group of high pleiotropy genes contained 38 WGD genes, significantly more than the 18 classified as low pleiotropy (p < 4.8e-3; Figure 6). TS-derived groups, from an overall smaller dataset, showed the same trend with 18 and 4 WGD duplicates in the high and low groups, respectively.

WGD gene pairs have been shown to sometimes have unequal allocation of importance, though they typically retain similar function (Kellis et al., 2004; VanderSluis et al., 2010), and pleiotropy roles reflect both these scenarios. We investigated the behavior of the duplicate partners of the DMA-derived high pleiotropy genes (most partners of genes in the TSA-derived pleiotropy groups have not been screened in SGA). First, considering only the handful of WGD pairs whose members both meet our degree criteria and therefore both have assigned pleiotropy classes, we find similarity in pleiotropy (Figure 6). Two WGD pairs were composed of two high-pleiotropy genes each (the pair ACE2 and SWI5, and the pair RPL40A and RPL40B) and, similarly, one pair had two low-pleiotropy members. However, no pairs of genes that both had high degree contained a low- and high-pleiotropy gene—a hint that duplicate partners of high pleiotropy genes tend to have higher pleiotropy than partners of low-pleiotropy genes (ranksum p < 0.016). Most partners of the genes with classified pleiotropy had lower degree. Of the 38 high-pleiotropy WGD genes, 28 duplicate partners have been screened as SGA queries and they show variation in both GI degree and pleiotropy scores (Figure 6).

**Figure 6.**
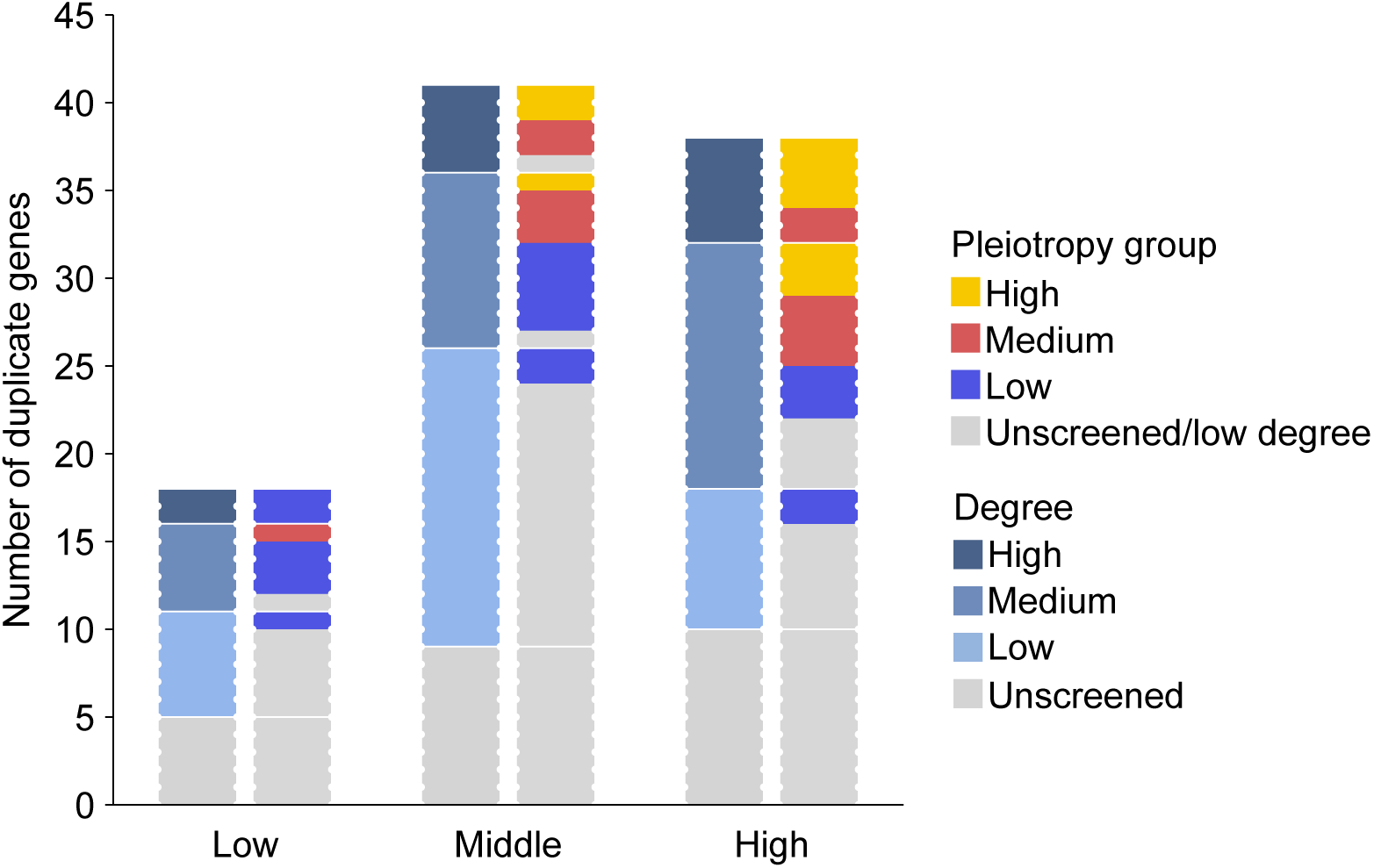
High-entropy genes are more likely to be whole-genome duplicates than are low-entropy genes. Bar heights show the number of whole-genome duplicate genes that are in the classes of high-, medium-, and low-pleiotropy, as labeled on the x-axis. GI degree and pleiotropy data shown as bar coloring describes the WGD *partners* of the high-degree classified genes. Bars on the left side show partner GI degree, where genes are considered “high” if their degree is at least the 60 percentile “hub” threshold used to define pleiotropy classes (near 100); “medium” if degree is at least 50 but lower than high cutoff; and “low” if degree is lower than 50. Bars on the right side show partner pleiotropy scores, which use the same thresholds as the standard pleiotropy classes defined for high-degree genes. Stacked sections of pleiotropy scores correspond to the matching sections of degree, as shown by horizontal white lines. For example, of all the WGD partners of classified low-pleiotropy genes, there are six with low GI degree (light blue, left bar) and, of these, one participated in enough biclusters that it could be given and pleiotropy score, which was low (one unit of dark blue, right bar). As another example, there are six high-pleiotropy genes whose WGD partners have high degree and, of these, four also have high pleiotropy. High-degree genes will be counted as both a classified gene and a partner of a classified gene.

A third copy-number based feature, copy number volatility (Wapinski et al., 2007), was also higher in high pleiotropy genes (p < 8.4e-4, Figure 4 A,B). This property measures the number of times a gene is lost or duplicated within extant or ancestral yeast species. We note that although copy number and copy number volatility are correlated with the binary WGD duplicate feature, these features each remain significantly associated with high pleiotropy genes after controlling for WGD duplication (p < 0.032 and < 0.011, respectively).

### Domains are more common in high-pleiotropy genes

The proteins of high pleiotropy genes tended to have more domains than those of low pleiotropy genes, a relationship supported by a recent GO-based measure of multifunctionality (Pritykin et al., 2015). Speculating that the association between the number of domains and pleiotropy is driven by functions of individual domains, we tested for enrichment of specific domains and combinations of domains, but did not find significant results for either medium- or high-pleiotropy genes.

### Characteristics of low pleiotropy genes

Genes that have low pleiotropy are characterized as highly prominent genes that are well-studied, conserved, and important. We found that low-pleiotropy genes are involved in more complexes than high pleiotropy genes (DMA p < 1.1e-5, TSA 7.5e-7), a characteristic derived from a literature-curated protein complex standard (Costanzo et al., 2016). For TSA-derived pleiotropy, this result, is also supported by a tendency to have a higher number of protein interactions in Tap-MS experiments (p < 6e-5) (Table 1). Participation of low-pleiotropy genes in protein complexes likely has the result of constraining the evolution of these genes (Lovell and Robertson, 2010).

A second characteristic of low-pleiotropy genes is that, compared to the high-pleiotropy genes, they have higher phenotypic capacitance, which is a measure of average morphological variance upon deletion of a nonessential gene and therefore also an indication of ability to buffer variability in phenotypes (Levy and Siegal, 2008). The authors who investigated phenotypic capacitors described a subset of capacitors that function in protein interaction clusters containing multiple capacitors and have a number of specific GO enrichments. This suggests that some capacitors promote phenotypic robustness by working in specific pathways. The abundance of these capacitors in our low-pleiotropy shows that our process of measuring a gene’s functional behavior through GI modules has distinguished between genes with specific roles in varied pathways (high pleiotropy) and genes whose deletion effects, but not necessarily wild-type behavior, has a variety of phenotypes.

Low pleiotropy genes have a higher number of annotations in the form of curated phenotypes (DMA p < 2.8e-4, TSA p < 2.1e-3) and GO annotations (TSA p < 8.6e-4). While at first this seems to contradict their status as focused functional influence, the annotation results likely belie investigation bias in our understanding of yeast. Indeed, for the TSA-derived pleiotropy groups, we find a tendency of the low-pleiotropy gene class to be old and conserved and to have a strong phenotype (Table 1, “Yeast conservation”, “Broad conservation”, “Age”, “Single mutant fitness defect)—exactly the type of genes that has attracted the most attention over the decades. The low-pleiotropy genes from the DMA-derived scoring configuration “query, adjacent” may also trend in the conserved direction, but it is not robust, with only two variations displaying it. Therefore, these more traditional measures of importance are not associated with high-pleiotropy genes. This suggests that our unbiased measure of pleiotropy captures an as-yet unappreciated amount of functional influence that flourishes in many newly evolved genes.

### Characteristics significantly associated with both high- and low-pleiotropy genes

All results discussed in previous sections were robustly associated with only one of the high and low pleiotropy classes, indicating agreement across the different scoring configurations we explored. However, we find that the features SMF and essentiality are significantly associated with both high- and low-pleiotropy genes, depending on the scoring configuration used (Table 1).

## Discussion

Genetic interactions provide a valuable view of pleiotropy by revealing gene functions at a molecular level. Clusters of within-pathway interactions highlight modules of genes related to specific cellular processes, like pathways or protein complexes, while between-pathway interactions occur when two pathways buffer each other. With this sensitivity to such a variety of gene-gene relationships, genetic interactions are well-suited for identifying diverse functions. Importantly, genetic interactions are calculated solely from phenotypic measurements, namely growth rates, in our case. Therefore, all genetic interactions represent functions that are evolutionarily relevant. Despite the fact that only one phenotype is measured, the functions represented by genetic interactions span most aspects of cellular biology (Costanzo et al., 2010; Costanzo et al., 2016).

Another strength of genetic interactions is their ability to reveal functions that may be undetectable in single mutants. A negative genetic interaction between two genes indicates a shared function that either one can perform individually, i.e. a buffering relationship. By assessing a gene’s pleiotropy within the GI network, we leverage the context of many (individual) background mutations, effectively removing layers of buffering and exposing the gene’s formerly hidden phenotypes. Some pleiotropy studies suggest that most genes affect few traits (refs for L-shaped distributions & “modular pleiotropy” model), but none of the considered datasets measure gene roles that are normally buffered in single mutants, leaving both theoretical and empirical discussions (Wang et al., 2010) to possibly underestimate pleiotropy. Still the importance of recognizing buffered functions depends on the extent to which individuals in natural populations harbor genetic variations that have genetic interactions.

A key element or our pleiotropy measure is the organization of the GI network into biclusters, which has multiple benefits. First, we have higher confidence in structures of genetic interactions than in individual interactions because dense clusters are very unlikely to occur by chance. Second, the functional level of a module removes redundancy by treating a set of genes as a unit. Because our method uses the associate-side of biclusters to determine annotations, genes that share a function are treated as a single unit. These functional units are summarized by an entropy measurement, the final pleiotropy score, which describes the shape of the distribution of modules among functions and differentiates broad from focused functional influence of a gene.

Through characterization of genes classified as having high and low pleiotropy, we found that evolution-related properties distinguished the groups. High-pleiotropy genes were more likely to be duplicated and to change in copy number throughout 30 yeast species. Contrasts in functional behaviors of the pleiotropy classes showed that high-pleiotropy genes have greater variability in expression, while low-pleiotropy genes are likely to be part of protein complexes. These interesting characterizations may shed light on the evolutionary processes through which genes may acquire multiple functions.

We propose that functional freedom is an important property enabling pleiotropy. Gene duplication and divergence is considered to be the primary source of raw material through which adaptions appear. The fact that duplicated genes tend to have high pleiotropy suggests that this process of new adaptations allows the accumulation of diverse functions in single genes, as opposed to only yielding two specialized (i.e. low-pleiotropy) genes. Partial functional buffering, relatively common between duplicates (Musso et al., 2008; VanderSluis et al., 2010), likely plays a role in this process. One model consistent with these ideas is subfunctionalization, in which the functions of the original/ancestral gene are partitioned between the paired duplicates. In this case, the two genes can complement each other such that each gene has functional regions maintained by selective constraint and regions that may mutate in a way that compromises the original function, but possibly acquire new functions. Even duplicates that are asymmetric in GI degree have been shown to maintain buffering relationships (VanderSluis et al., 2010). A second mechanism by which duplicates may diverge and become pleiotropic is suggested by the tendency of high-pleiotropy genes to have high variance in expression. Changes in regulatory patterns occurring soon after duplication may provide a route for acquiring environment-specific roles. Functions unneeded in particular conditions may be altered to respond to new challenges, thus diversifying the gene’s functions. Acquisition of new functions through variable expression is not limited to duplicates, but is proposed as a general mechanism promoting adaptation to stress environments. Finally, the significantly low number of pleiotropic genes that have membership in protein complexes suggests an avoidance of evolutionary constraint of sequence changes and consequent barrier to gaining novel functions.

While the characterization of pleiotropic genes as being sheltered from functional constraints provided by duplicates buffering each other and as lacking physical interactions in protein complexes offers insight into the kind of genes that are able to acquire new functions, it remains to be shown how pleiotropic genes have risen to such prominence that they are genetic interaction hubs. Indeed, the functional freedom suggested by our characterization of pleiotropic genes is a contrast to Fisher’s classic geometric model of pleiotropy, which predicts that pleiotropic genes will be evolutionarily constrained and has been advanced by the “cost of complexity” model (Orr, 2000; Welch et al., 2003). However these characterizations can coexist at different time periods in a gene’s life cycle: pleiotropy may originate over a relatively short period of time following de novo birth of a gene, a gene duplication, or a regulatory change buffered by a non-duplicate alternative pathway, and subsequent loss of buffering. Intriguingly, the TSA-derived high-pleiotropy genes contained a significantly higher number of Saccharomyces-specific genes as compared to the low-pleiotropy genes, which is evidence that participation in many processes can occur near the beginning of the lifecycle of a gene. Additionally, there may be mechanisms in place that stabilize the effects of evolving genes. Post-transcriptional regulation may be strong enough to counteract expression-level patterns, therefore stabilizing protein levels when needed (Artieri and Fraser, 2014), and explaining the association between high-pleiotropy genes and high protein abundance. Similarly, genetic hubs have been shown to typically have steady expression levels, likely as a consequence of their importance (Park and Lehner, 2013), but the hubs that have variable expression levels are enriched for duplicates that may be able to buffer the effects of low expression (Park and Lehner, 2013), meaning duplication is not only a source of novel adaptations, but may also be a mechanism by which networks are robust.

## Acknowledgements

This work was partially supported by the National Institutes of Health (R01HG005084, R01HG005853) and the National Science Foundation (DBI 0953881). BA, CB, and CLM are also fellows of the Canadian Institute for Advanced Research (CIFAR) Genetic Networks Program. Computing resources and data storage services were partially provided by the Minnesota Supercomputing Institute and the UMN Office of Information Technology, respectively. EK was partially supported by an NSF Graduate Research Fellowship and a U. of Minnesota Doctoral Dissertation Fellowship.

## Methods

### XMOD overview

Biclusters were generated from each network according to the procedure described by Bellay et al. (2011). In this method, frequent itemset mining is used to find groups of array genes that frequently (with high support) occur together as subsets of query genes’ interactors. To determine statistical significance, the biclusters are compared to biclusters mined from ten randomized versions of the network. All biclusters are assigned a score that represents the likelihood of all their contained interactions occurring if genes interacted randomly, conditioned on the genes’ interaction degrees. The scores of the random biclusters are used as a null distribution to assign p-values to the real biclusters, which are expected to have lower scores due to non-random gene associations. Biclusters with p-values higher than a chosen significance level are discarded.

The nature of our new SGA data motivated a number of modifications and additions to this bicluster-discovery framework, which are detailed in the following paragraphs.

### Binary networks input to mining

The TSA (temperature sensitive array) and DMA (deletion mutant array) SGA GI networks from Costanzo et al. (2016) were binarized by defining interacting strains as pairs with epsilon scores less than or equal to −0.08, according to the established intermediate cutoff. We additionally added self interactions to the network, to allow the discovery of cliques.

The set of queries in each SGA network includes mutant strains with DaMP and TS alleles of essential genes, often with multiple alleles of a single gene. Due to the biased multiplicity of many essential genes within the set of query strains, bicluster datasets generated from the complete data set may be uninteresting or difficult to interpret because bicluster significance would be driven by the highly correlated behavior of alleles of the same gene. To overcome this problem, we produced 15 replicates of each network, each containing one randomly selected allele for each gene. Using many replicate networks yields good representation of different alleles and allows different combinations of alleles to be chosen. In all, 15 replicates of each of the DMA and TSA GI network were input to separate runs of XMOD.

### Bicluster discovery

All frequent item set mining, done within the XMOD framework, was performed using an implementation of the Eclat algorithm (Zaki et al., 1997) by Christian Borgelt, which is available at http://www.borgelt.net/eclat.html, using the “-tc” option to report only closed item sets. A single test run on the DMA network yielded over 37 million biclusters with a size of at least four query strains and four array strains. Based on the observation that only ∼2.7% of all 4x4, 4x5, and 5x4 biclusters are significant at a p-value threshold of 10e-4, we used the Eclat option to remove from the results all biclusters of these three mentioned sizes and those with either dimension smaller than four in order to reduce memory usage in XMOD; the Eclat options to accomplish this are “-s-4 –m-4-F-6-5-4”. After eliminating these small biclusters, the DMA network replicates contained between an average of ∼24.5 million biclusters and the TSA network replicates contained an average of ∼20 million biclusters.

As described in Bellay et al. (2011) and briefly above, XMOD determines empirical p-values of biclusters through comparison to biclusters discovered in ten randomized networks. Combining biclusters from ten random networks supplies a better sampling of biclusters of larger dimensions than one random network could, however, there was an overabundance of random-network biclusters with small dimensions (e.g. 4x6, 6x4, 5x5). So for better speed and memory use, we randomly discarded biclusters to keep a maximum of two million for each size.

All biclusters with p-values >= 1e-4 we discarded, leaving ∼14m (57%) DMA and ∼10m (50%) TSA biclusters per network replicate. The vast majority of biclusters containing 30 or more interactions (e.g. 6x6, 7x5, and larger) were significant, since very few large biclusters were found in the randomized networks.

### Preference of bicluster sizes during condensing

Bicluster discovery through frequent item set mining typically produces modules that overlap, i.e. a bicluster usually shares some of its interactions with other biclusters. While this certainly reflects reuse of genes in different cellular functions, it is also caused by our inability to discover larger modules that are fractured by false negatives (biological or technical). We therefore used a method described in Bellay et al. (2011) that condenses a set of biclusters by identifying pairs of overlapping biclusters and removing one, leaving a non-redundant set of modules. The procedure is greedy and proceeds as follows: First, order all biclusters from best to worst. Then, select the biclusters in order and upon the selection of a bicluster, remove overlapping biclusters from any future consideration. We defined “overlapping” as the smaller bicluster having 10% or more genetic interactions in common with the larger.

To define the best-to-worst ordering, we determined preferences for different bicluster sizes, and built a size-lookup table to pick between differently sized biclusters. For our use, the quality of a bicluster can be measured by how well it reiterates a set of genes annotated by a GO term. We selected Jaccard similarity between the bicluster gene set and an enriched GO term as a simple statistic to measure this. Since calculating GO term enrichments on all bicluster gene sets would take too long, we used a sample of biclusters to rank bicluster sizes (expressed in two dimensions) according to results from Jaccard similarity analysis. First, for each bicluster size, we collected all biclusters up to a maximum of 10,000 and removed redundancy from this set by consecutively selecting biclusters in random order and removing any other bicluster from future selection if more than 10% of its interactions overlapped with the selected one. Next, statistically enriched GO terms were determined for every bicluster (using genes from one side) and the maximum Jaccard similarity obtained from each bicluster was recorded, yielding a distribution of maximum Jaccard similarities for each bicluster size. Finally, as a summary statistic representing likelihood of reflecting a known module, we kept the median Jaccard score for each bicluster size, organized as a lookup table to consult. This analysis was done separately for the DMA- and TSA-derived biclusters.

A size preference table is specific to the bicluster dataset (either TSA or DMA) and the side of the bicluster (query-strain or array-strain) that is intended to be used in further analyses. Therefore, a total of four tables were created (supplemental file).

### Bicluster functional profiles

For both the TSA and DMA network, every strain has 15 sets of biclusters that it appeared in, one set from each of the network replicates. All of these were condensed individually (i.e. there were 15 different sets of biclusters per strain, per network) twice: first for the purpose of annotating the query-strain sides of biclusters, and second for the purpose of annotating the array-strain sides, using the appropriate size preference table in each case. The condensed set of biclusters intended for query-side annotations was used, for example, in the “Query, associate” scoring configuration, while the array-side set was used for the “Query, adjacent” scoring configuration.

To create functional profiles from the sets of condensed biclusters, we annotated one side of each bicluster with biological process terms that have been manually annotated (MA, (Costanzo et al., 2010), supplementary file) or systematically annotated (SAFE, (Baryshnikova, 2016; Costanzo et al., 2016), supplementary file) to yeast genes. Every bicluster was annotated by any term for which its gene set had significant enrichment or to which at least 40% of queries were annotated. The numbers of annotations to each term were counted and normalized within each of the 15 sets of biclusters associated with each query or array strain, creating functional profiles. These replicate profiles were averaged, yielding one bicluster-based functional profile per strain.

### Pleiotropy (entropy) scores

Entropy of a strain was calculated from its functional profile as 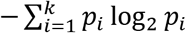, where *p_i_* is the fraction of biclusters annotated with the *i*-th process and *k* is the total number of terms in the annotation scheme. Pleiotropy scores were not assigned to genes that had fewer than 10 biclusters.

### Validation of profiles

To compare each gene’s bicluster-derived annotation profile to its gold standard annotations, either MA or SAFE, we used a simple one-dimensional version of the clustering algorithm DBScan (Ester et al., 1996) to find the most striking highly annotated process or processes for each functional profile. Before clustering, we normalize each profile by dividing by its maximum value. Our implementation of DBScan visits values from highest to lowest and labels a value as an outlier if it has no neighbors (minPts parameter is 1) at a distance of less than 0.2 (Eps parameter), otherwise it defines a cluster and expands the cluster following the standard algorithm. We defined profile-predicted annotation as (1) all outliers that are higher than the first cluster, if there are any, or (2) the highest cluster, if there are no outliers.

### Testing for associations between pleiotropy and gene characteristics

We used Wilcoxon ranksum tests to compare the values of gene characteristics of high-pleiotropy genes to those of low pleiotropy genes. The 37 physical, functional, and evolutionary gene properties are listed in the section “Gene properties.” We performed tests using pleiotropy scores obtained from the six different pleiotropy scoring configurations and testing variants, which are described below. The significance of p-values from ranksum tests was determined using the FDR-control procedure described in Benjamini et al. (2006), treating the sets of 37 tests with identical set-ups, but different gene properties, as families.

The scoring configurations comprise the following: network orientations described in the “Network inputs to mining” (query vs array), the bicluster side used for functional enrichment described in the section “Pleiotropy (entropy) scores” (associate vs adjacent), and the gene annotations used (MA vs SAFE). The six combinations of these methods choices that are used are “Query, associate, manual”, “Query, associate, SAFE”, “Query, adjacent, manual”, “Array, associate, manual”, “Array, associate, SAFE”, and “Array, adjacent, manual”.

For each of scoring configurations, we performed multiple ranksum tests, called “testing variants,” that explore different ways to define the high- and low-pleiotropy classes and control for possible biases that different types of mutant alleles may cause. The methods choices we considered that affect any type of gene are: controlling for the GI degree of genes (DC vs noDC); removing strains that show weak signs of batch effects (rmBadBatch); and altering the percent of genes that are added to the high and low pleiotropy classes, which is set to 30% by default (tail20, tail40). Variants related to genes represented by TS and DAmP alleles are the following: determining the gene’s pleiotropy score by taking the mean score or maximum score of the alleles (average vs max); removing one type of mutant allele (noDamp, noTs); applying a minimum GI degree threshold of 50 before averaging the degree of alleles to determine the high-degree genes that may be classified (damp50, ts50). The combinations of these methods choices that are used are described in Table SXX (“ranksumTestVariantsTable.xlsx”).

The pleiotropy classes of high, medium, and low, are determined for each pairing of scoring configurations and testing variants because these methods choices affect which genes are considered. First, high-degree genes are identified as those with degree in the top 40% out of all genes that have been screened, have a pleiotropy score, and have not been removed by one of the test-variant modifications. For test variants that include degree control, we regress pleiotropy scores against degree and keep the pleiotropy residuals in place of pleiotropy scores. Then the pleiotropy scores (or score residuals) are divided into classes with the genes whose pleiotropy is in the top 30% of the high-degree genes labeled high-pleiotropy, genes in the bottom 30% labeled as low-pleiotropy, and the remaining 40% of genes labeled as medium pleiotropy.

### Gene properties

**Age 1-20** indicates the phylogenetic distance of the most distantly related species with an identified ortholog to a given yeast gene. Genes only found in *S. cerevisiae* are assigned the age of 0 and genes with orthologs appearing in more distant species are assigned higher ages. Two phylogenetic trees were used in this analysis: one obtained from Ostlund et al. (2010) contains 100 animal, plant, and fungi species and one obtained from Wapinski et al. (2007) contains 23 yeast species.

**Broad conservation** is a count of how many non-yeast species, out of a set of 86, have an ortholog of a given gene. To count this, we obtained orthogroup designations from InParanoid (Ostlund et al., 2010). For each gene, we considered it to have an ortholog in another species only if it appeared in a cluster with the other species and was given a score of 1.0 by the InParanoid clustering method; that is, we considered a yeast gene to have an ortholog in species x if it was a seed gene for a gene cluster that had an orthologous cluster in species x. Note that this measure is similar, though complementary, to the “yeast conservation” measure described below, which focuses on conservation within the yeast clade.

**CAI**, codon adaptation index, is a sequence-based measure of bias in usage of synonymous codons as compared to usage in highly expressed genes. It was calculated using the cai tool and the default codon usage table in the EMBOSS suite (Rice et al., 2000).

**Chemical-genetic degree** is a count of drug and environmental conditions to which a homozygous diploid gene-deletion mutant strain is significantly sensitive (Hillenmeyer et al., 2008).

**Coexpression degree** is a measure derived from a co-expression network based on integration of a large collection of expression datasets (Huttenhower et al., 2006). The network was sparsified by considering only edges between gene pairs whose co-expression levels were above the 95^th^ percentile. The co-expression degree of a gene is the number of genes with which its co-expression value is retained in this restricted network.

**Complex member** is a binary feature that indicates whether the corresponding protein is a component of at least one complex based on the complex standard provided in Costanzo et al. (2016).

**Copy number** is a count of the number of paralogs each gene has. This was determined from clusters identified by the InParanoid algorithm (Ostlund et al., 2010). All genes that appeared in the same cluster were considered paralogs.

**Copy number volatility** is the number of times that a gene is lost or gained among 23 Ascomycete fungi species, as defined by Wapinski et al. (2007).

**Curated phenotypes** Mutant phenotypes were downloaded from the Saccharomyces Genome Database (SGD) on January 31, 2013. The list of phenotypes was filtered to include only those related to deletion mutants of verified or uncharacterized open reading frames (mutant type = ‘null’, feature type = ‘ORF’). Phenotypes were further filtered to only include increased or decreased phenotype expression compared to a wild-type strain. Finally, the number of non-wild-type phenotypes was counted for each gene. Unclear descriptions of phenotypes, such as “abnormal”, were ignored.

**Deleterious SNP rate** is the number of predicted deleterious SNPs observed for a given gene in the recently sequenced set of diverse S. cerevisiae strains (Liti et al., 2009) normalized by gene length. **Deleterious SNP rate of strains** is similar, but counts strains containing deleterious SNPs. These SNP features were derived from identification and analysis of SNPs in 19 strains as described in (Jelier et al., 2011). Briefly, SNPs were identified from sequence alignments of all strains to the S288C reference sequence. The SIFT algorithm, with some modifications, was used to predict which nonsynonymous SNPs are likely to have functional consequences. We applied the recommended threshold to SIFT scores, calling any SNP with a score of <= 0.05 deleterious.

**dN/dS** is the ratio of the number of nonsynonymous to synonymous mutations in a given gene. We computed the average dN/dS ratio for *S. cerevisiae* in comparison to the sensu strictu yeast species (*Saccharomyces paradoxus*, *Saccharomyces bayanus*, and *Saccharomyces mikatae*). Protein sequences were aligned using MUSCLE and dN/dS ratios were computed using PAML (Edgar, 2004; Yang, 2007).

**Effective number of codons** is a measure of codon usage bias and is an alternative to CAI that does not require a pre-defined set of highly expressed genes. This measure was computed using PAML.

**Essential**, a binary feature, is true for any gene that is required for viability under standard laboratory conditions.

**Expression level** is a measurement of the mRNA expression level of a gene (Holstege et al., 1998).

**Expression variance, environ.** is the variance in a gene’s expression across all measurements in the Gasch et al. (2000) dataset. This study subjected yeast to many evironmental conditions and measured expression of nearly all yeast genes with microarrays. Environments included heat shock, hydrogen peroxide, superoxide generated by menadione, diamide, dithiothreitol, hyper-osmotic shock, amino acid starvation, nitrogen source depletion, and progression into stationary phase, as well as alternative carbon sources and variable temperatures. The data contain multiple time points and temperatures for the environments listed.

#### Expression variance, genetic-A

The expression dataset produced by Brem and Kruglyak (2005) measures expression of 6162 genes in the strains BY4716, RM11-1a, and 112 segregants from crosses between BY4716 and RM11-1a. This expression variance feature is simply the variance across these strains, measured for each gene, and represents the amount of variation that occurs throughout genetically diverse genomic backgrounds.

#### Expression variance, genetic-B

Skelly et al. (2013) obtained 22 yeast strains from geographically and environmentally diverse locations and performed RNA-seq to measure gene expression levels. This expression variance feature is the variance across the strains, measured for each gene, and represents expression variation that occurs in genetically diverse genomic backgrounds.

**log2(Distance from telomere)** is the distance, in nucleotides and log-transformed, between a gene and the start of a its closest telomere.

**Multifunctionality** is a count of annotations to “biological process” terms of the Gene Ontology. Specifically, the total number of annotations across a set of functionally distinct GO terms described in Myers et al. (2006) was used as a multi-functionality index.

**Number of complexes** is a count of the number of complexes by which a given gene is annotated in the protein complex standard provided in Costanzo et al. (2016).

**Number of domains** is the number of domains, counting repeated domains, present within a given protein, as identified by PFAM (downloaded July 2015). **Number of unique domains** is the same but does not count repeated domains.

**Originated in Saccharomyces** is a binary value that is true if a gene originated in the Saccharomyces clade of the phylogenetic tree, which is assumed if the most distant species with an ortholog is a Saccharomyces yeast species. Specifically, we consulted the species tree from Wapinski et al. (2007) and identified all genes that appear only in *S. cerevisiae*’s closest relatives: species up to and including *Saccharomyces bayanus*. Note that although some more distant species (*Naumovozyma castellii*, *Lachancea kluyveri*) were originally placed in the genus *Saccharomyces* and may still be referred to with this name as described in Wapinski et al., these have subsequently been associated with different genera.

**Phenotypic capacitance** was computed by Levy and Siegal (2008) and captures variability across a range of morphological phenotypes upon deletion of each of the nonessential genes.

#### Pleiotropy sum

##### PPI degree, Tap MS

Physical interaction degree from Tandem Affinity Purification coupled with Mass Spectrometry (TAP-MS) refers to the total number of interactions in the union of the Gavin et al. (2006) and Krogan et al. (2006) datasets.

**PPI degree, Y2H** is the total number of binary, physical interactions detected using yeast two-hybrid analysis (Yu et al., 2008).

**Protein abundance** was measured by fluorescence of GFP-tagged proteins grown in liquid rich media; **protein abundance under stress** was measured by fluorescence of GFP-tagged proteins grown in liquid minimal media (Newman et al., 2006).

**Protein disorder** is the percent of unstructured residues as predicted by the Disopred2 software (Ward et al., 2004).

**Protein length** is the number of amino acids in a gene’s associated protein.

**Single mutant fitness defect** was calculated by Costanzo et al. (2016).

**SSD duplicate**, a binary feature, is true for genes with one or more paralogs that resulted from small scale duplication (SSD) events. To identify pairs of genes that emerged from SSD events, we searched for gene pairs that meet the following criteria: the gene pair must have a sufficiently high sequence similarity score (FASTA Blast, E = 10), sufficient protein alignment length (> 80% of the longer protein), an amino acid level identity of at least 30% for proteins with aligned regions longer than 150 amino acids or greater than [0.01n + 4.8L^(-0.32(1 + exp(-L/1000)))] with L defined as the aligned length and n = 6 for shorter proteins (Gu et al., 2002; Rost, 1999).

#### Transcription level

**WGD duplicate**, a binary feature, is true for any gene that has a paralog that resulted from the whole genome duplication event. The WGD event designation was reconciled from several sources (Byrne and Wolfe, 2005).

**Yeast conservation** counts how many of 23 different species of Ascomycota fungi possess an ortholog of a gene. This measure was described by Wapinski et al. (2007) and ortholog data were downloaded from the associated website http://www.broadinstitute.org/regev/orthogroups/.

